# Evolutionary history of cotranscriptional editing in the paramyxoviral phosphoprotein gene

**DOI:** 10.1101/2020.09.30.321489

**Authors:** Jordan Douglas, Alexei J. Drummond, Richard L. Kingston

## Abstract

The phosphoprotein gene of the paramyxoviruses encodes multiple protein products. The P, V, and W proteins are generated by transcriptional slippage. This process results in the insertion of non-templated guanosine nucleosides into the mRNA at a conserved edit site. The P protein is an essential component of the viral RNA polymerase, and is encoded by a direct copy of the gene in the majority of paramyxoviruses. However, in some cases the non-essential V protein is encoded by default and guanosines must be inserted into the mRNA in order to encode P. The number of guanosines inserted can be described by a probability distribution which varies between viruses. In this article we review the nature of these distributions, which can be inferred from mRNA sequencing data, and reconstruct the evolutionary history of cotranscriptional editing in the paramyxovirus family. Our model suggests that, throughout known history of the family, the system has switched from a P default to a V default mode four times; complete loss of the editing system has occurred twice, the canonical zinc finger domain of the V protein has been deleted or heavily mutated a further two times, and the W protein has independently evolved a novel function three times. Finally, we review the physical mechanisms of cotranscriptional editing via slippage of the viral RNA polymerase.

## 1 Introduction

The *Paramyxoviridae* are a family of nonsegmented, negative-sense, single-stranded RNA viruses, within the order Mononegavirales (Amarasinghe et al., 2019; Pringle, 1991; Rima et al., 2018). Type species include measles virus (MeV; genus: Morbillivirus), mumps virus (MuV; genus: *Orthorubulavirus*), Sendai virus (SeV; genus: *Respirovirus*), and Hendra virus (HeV; genus: *Henipavirus*). The *Paramyxoviridae* appear to infect most vertebrate species and are responsible for a number of serious diseases in both animals and humans.

The single-stranded RNA genome is bound to the viral nucleocapsid protein, forming a helical protein-nucleic acid complex which organises and protects the genome. The nucleocapsid acts as a template for both gene transcription and genome replication, processes that are executed by the viral RNA-dependent RNA-polymerase (RdRp; Whelan et al. (2004); Noton and Fearns (2015); Fearns and Plemper (2017); Guseva et al. (2019)). Each nucleocapsid protein binds six nucleotides of RNA (Gutsche et al., 2015; Alayyoubi et al., 2015; Jamin and Yabukarski, 2017; Webby et al., 2019), and paramyxoviral genomes always conform to the “rule of six” whereby genome length is some multiple of six (Kolakofsky et al., 1998; Calain and Roux, 1993; Kolakofsky et al., 2005). This is hypothesised to result from the requirement to position the promoter sequences required for initiation of RNA synthesis in the correct register, or phase, with respect to the nucleocapsid protein.

The phosphoprotein (P protein) is the non-catalytic subunit of the viral RdRP and mediates interaction between the polymerase and the nucleocapsid. The P protein is essential for translocation of the RdRP along its template, among other roles (Kingston et al., 2004; Du Pont et al., 2019; Bruhn et al., 2019; Milles et al., 2018; Guseva et al., 2019). The P gene, however, encodes multiple protein products through a combination of non-canonical transcription and translation events. This phenomenon, known as overprinting, is particularly common in viruses as it increases the utility of the genome, which is often under strong size constraints (Brandes and Linial, 2016; Sabath et al., 2012).

Cotranscriptional editing at the P gene governs production of the P, V, and W proteins. Production of multiple proteins results from insertion of one or more non-templated guanosine nucleosides (G) into the mRNA at a conserved edit site, through transcriptional slippage (Vidal et al., 1990b; Hausmann et al., 1999a). This process of base insertion stochastically shifts the reading frame. As a result, the P, V, and W proteins share a common N-terminal region (encoded by the gene sequence upstream of the edit site), but possess distinct C-terminal regions (encoded by the gene sequence downstream of the edit site), conferring differing functions on these three proteins. The mRNA encoding P, V, and W occur at different frequencies in different paramyxoviruses.

Although not the focus of this paper, operations at the translational level (Firth and Brierley, 2012), including leaky scanning (Giorgi et al., 1983; Shaffer et al., 2003), non-AUG initiation (Boeck et al., 1992; Curran and Kolakofsky, 1988), and ribosomal shunting (Latorre et al., 1998), facilitate production of yet more proteins from the P gene in many paramyxoviruses.

In this article we review cotranscriptional editing of the P gene in the *Paramyxoviridae*. We briefly discuss the assigned functions of the P, V, and W proteins, collate the available experimental information on the size distribution of guanosine nucleoside insertions, and propose a maximum parsimony model for evolution of the editing system. We also review what is known about transcriptional slippage and its connection with the genomic sequence at the edit site and displacement of nucleocapsid proteins from the genome.

## 2 Cotranscriptional editing in the *Paramyxoviridae*

Cotranscriptional editing at the P gene occurs through the insertion (or in certain mutants the deletion; Jacques et al. (1994)) of *m ∈* {0, 1, 2, …} guanosines, *G*_*m*_, into the mRNA at a conserved edit site. A *G*_3*k*+1_ insertion (*m* = 1, 4, 7, …) shifts the reading frame downstream of the edit site by −1 (or alternatively +2). A *G*_3*k*+2_ nucleotide insertion (*m* = 2, 5, 8, …) shifts the reading frame by −2 (or alternatively +1). A *G*_3*k*_ insertion (*m* = 0, 3, 6, …) leaves the reading frame unaltered.

Proteins translated from a *G*_3*k*+*n*_ mRNA species are considered to be functionally equivalent across all values of *k ∈*{0, 1, 2,} The mRNA flanking the edit site encodes an intrinsically disordered region of the protein (Habchi and Longhi, 2012; Guseva et al., 2019). Any extended sequence of G nucleotides is translated into polyglycine. While the conformational preferences of polyglycine are still not entirely established (Tran et al., 2008; Ohnishi et al., 2006), the homo-polymeric sequence will be disordered, and small variations in the length of this sequence are likely to be functionally neutral in this context.

The cotranscriptional editing system for the P gene operates in two different modes (**Figure 1**). In the 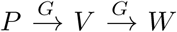 mode, P is encoded by the unedited gene. This is the situation in MeV (Cattaneo et al., 1989) and SeV (Vidal et al., 1990a). In the 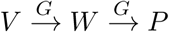 mode, V is the default and a *G*_3*k*+2_ insertion is required to encode P. This is the situation in MuV (Paterson and Lamb, 1990).

**Figure 1:**
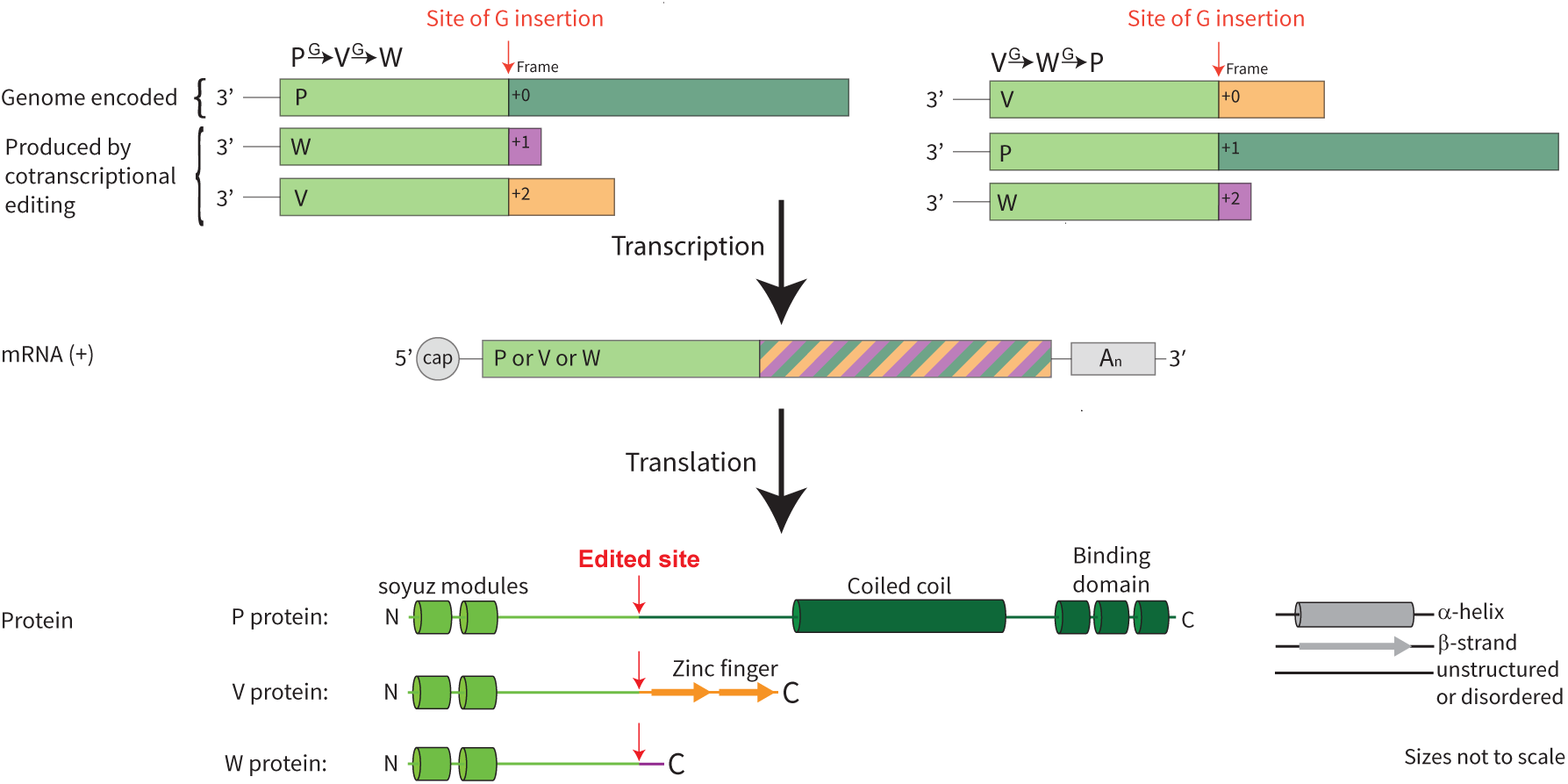
Cotranscriptional editing of the P gene. The two observed modes of editing are depicted: these are the 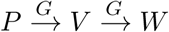 and 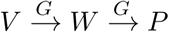 modes. A single transcript can encode one of P, V, or W depending on the number of guanosines stochastically inserted at the edit site during transcription. While the P, V, and W proteins share a common N-terminal region, their C-terminal regions are distinct.

We note that switching between these two edit modes requires a frameshift mutation in the genome, i.e. during genome replication. This mutation must occur at a position upstream of the edit site, but not so far upstream that it disrupts some other function of the encoded P protein. Moreover, due to the rule of six, any insertion or deletion (indel) must be rapidly compensated such that the genome length remains divisible by six. Otherwise the replication efficiency of the virus would be severely impacted (Calain and Roux, 1993; Kolakofsky et al., 2005; Skiadopoulos et al., 2003; Sauder et al., 2016). For example, a single nucleotide insertion upstream and proximal to the edit site, accompanied by a single nucleotide deletion elsewhere in the genome, would be sufficient to transit the system from 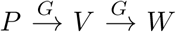 to 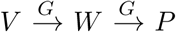.

### 2.1 P protein

The phosphoprotein has a range of functions. In complex with the viral large protein (L protein) it forms an integral part of RdRP. However, due to its greater relative abundance – a result of the “transcriptional gradient” that exists in all paramyxoviruses (Cattaneo et al., 1987; King et al., 2011) – it must also function independently of the L protein. The disordered N-terminal region of P is shared with V and W and contains the highly conserved soyuz1 and soyuz2 motifs (Karlin and Belshaw, 2012). These modules, together with internally located sequences, are involved in chaperoning the viral nucleocapsid protein monomers during replication (Yabukarski et al., 2014; Milles et al., 2018; Alayyoubi et al., 2015; Guryanov et al., 2016). This region also binds host proteins such as the STATs (Li et al., 2019; Devaux et al., 2007; Röthlisberger et al., 2010; Puri et al., 2009; Ciancanelli et al., 2009; Devaux et al., 2010; Chinnakannan et al., 2014), and its function is likely regulated by phosphorylation (Saikia et al., 2008; Sun et al., 2009; Sugai et al., 2012; Young et al., 2019; Pickar et al., 2014). The N-terminal region ranges from 109 aa (in APMV-3) to 570 aa (in GH-M74a) in length.

Downstream of the edit site, the unique region of the phosphoprotein contains an oligomerisation domain (a coiled coil; Burmeister et al. (2000); Tarbouriech et al. (2000); Communie et al. (2013a); Cox et al. (2013); Bruhn et al. (2014)) and a nucleocapsid / large binding domain (the foot domain, or X domain; Johansson et al. (2003); Kingston et al. (2008); Yegambaram et al. (2013); Blanchard et al. (2004)) which are connected by a flexible linker. The C-terminal region of the phosphoprotein binds to both the large protein (Bruhn et al., 2019; Abdella et al., 2020) and the nucleocapsid (Kingston et al., 2004; Habchi et al., 2011; Communie et al., 2013b; Bloyet et al., 2016; Du Pont et al., 2019) and mediates their engagement. The phosphoprotein is therefore essential (Curran et al., 1991) and is encoded by all paramyxoviruses. P is the largest of the three P gene proteins. C-terminal regions range from 229 aa (in PIV-5) to 386 aa (in CPIV-3).

### 2.2 V protein

The V protein regulates viral genome replication (Horikami et al., 1996; Witko et al., 2006; Parks et al., 2006; Nishio et al., 2008; Sleeman et al., 2008; Yang et al., 2015) and interferes with the innate immune response to viral infection (see reviews: Fontana et al. (2008b); Ramachandran and Horvath (2009)). The latter is linked to increased viral virulence (Devaux et al., 2008; Alamares et al., 2010; Schaap-Nutt et al., 2010). V has been reported to bind a multitude of proteins involved in activation of the type I interferon response, sometimes in a genus specific fashion. These proteins include cytoplasmic pattern recognition receptors RIGI, MDA5 (Motz et al., 2013), and LGP2 (Ramachandran and Horvath, 2010; Mandhana et al., 2018; Sanchez-Aparicio et al., 2018; Andrejeva et al., 2004; Childs et al., 2007; Rodriguez and Horvath, 2014), as well as TRIM25 (Sanchez-Aparicio et al., 2018), PP1 (Davis et al., 2014), MAVS (Sun et al., 2019), PLK1 (Ludlow et al., 2008), UBXN1 (Uchida et al., 2018), DDB1 (Li et al., 2006a; Salladini et al., 2017), various interferon regulatory factors (Takeuchi et al., 2003; Kitagawa et al., 2013; Palosaari et al., 2003; Lu et al., 2008), NF-*κ*B (Schuhmann et al., 2011), and the STAT proteins (Li et al., 2019; Didcock et al., 1999; Parisien et al., 2002). It has also been reported to interact with interferon stimulated gene products such as tetherin (Ohta et al., 2017, 2016) and other host proteins such as PKB/AKT1 (Sun et al., 2008).

The unique C-terminal region of V contains a highly conserved cysteine-rich zinc finger domain, which binds two zinc ions (Li et al., 2006a; Motz et al., 2013). The functions conferred by this unique C-terminal region are not always readily decoupled from the activities of the common N-terminal region. For instance, both of these regions bind STAT proteins (Li et al., 2019), and the structurally characterised interaction with DDB1 (Li et al., 2006a) is mediated by sequences from both the common N-terminal region and the unique C-terminal region of V.

V is the second largest of the P gene proteins: C-terminal regions range from 50 aa (in NiV) to 188 aa (in CPIV-3). While V aids viral replication, it is non-essential and is encoded by most but not all paramyxoviruses (Section 3.2).

### 2.3 W protein

A third protein may also be generated by contranscriptional editing. Unlike P and V, its unique C-terminal sequence is not conserved across, or even within, paramyxoviral genera and consequently this protein has been assigned many names (Fontana et al., 2008a) including W (Vidal et al., 1990a), D (Pelet et al., 1991; Galinski et al., 1992), PD (Wells and Malur, 2008), and I (Paterson and Lamb, 1990). For the purposes of this review, we use ‘W’ to denote the protein encoded by the reading frame that encodes neither P nor V. There is evidence that W has evolved a function within some paramyxoviral genera, though this is not always the case.

In Newcastle disease virus (NDV; genus: *Orthoavulavirus*), Hendra and Nipah virus (HeV and NiV; genus: *Henipavirus*), and Human parainfluenza virus 3 (HPIV-3; genus: *Respirovirus*), W accumulates in the nucleus (Karsunke et al., 2019; Yang et al., 2019; Shaw et al., 2005; Lo et al., 2009; Wells and Malur, 2008). Nuclear localisation signals can be identified in the unique region of the W protein (Shaw et al., 2005; Karsunke et al., 2019; Audsley et al., 2016a; Wells and Malur, 2008; Smith et al., 2018).

NDV sits alone, and we could not detect a homologous C-terminal region in the W protein of any other *Orthoavulaviruses*. A recent study showed that deleting the C-terminus of the W protein impaired NDV replication in cultured cells, and this effect was relieved when the full-length W protein was supplied in trans (Yang et al., 2019). However, no detailed function has been assigned to this protein.

The *Henipavirus* W protein influences the course of disease in animal models (Satterfield et al., 2015, 2016), and may play a direct role in subversion of the type I IFN response (Ciancanelli et al., 2009; Shaw et al., 2005). This protein also modulates host gene expression by interacting with the 14-3-3 family of regulatory proteins, an interaction that depends upon phosphorylation of the penultimate serine residue (Edwards et al. (2020); **Fig. 2**). However as with the V protein, it has proved difficult to experimentally decouple common functions conferred by the N-terminal region, with unique functions conferred by the C-terminal region.

**Figure 2:**
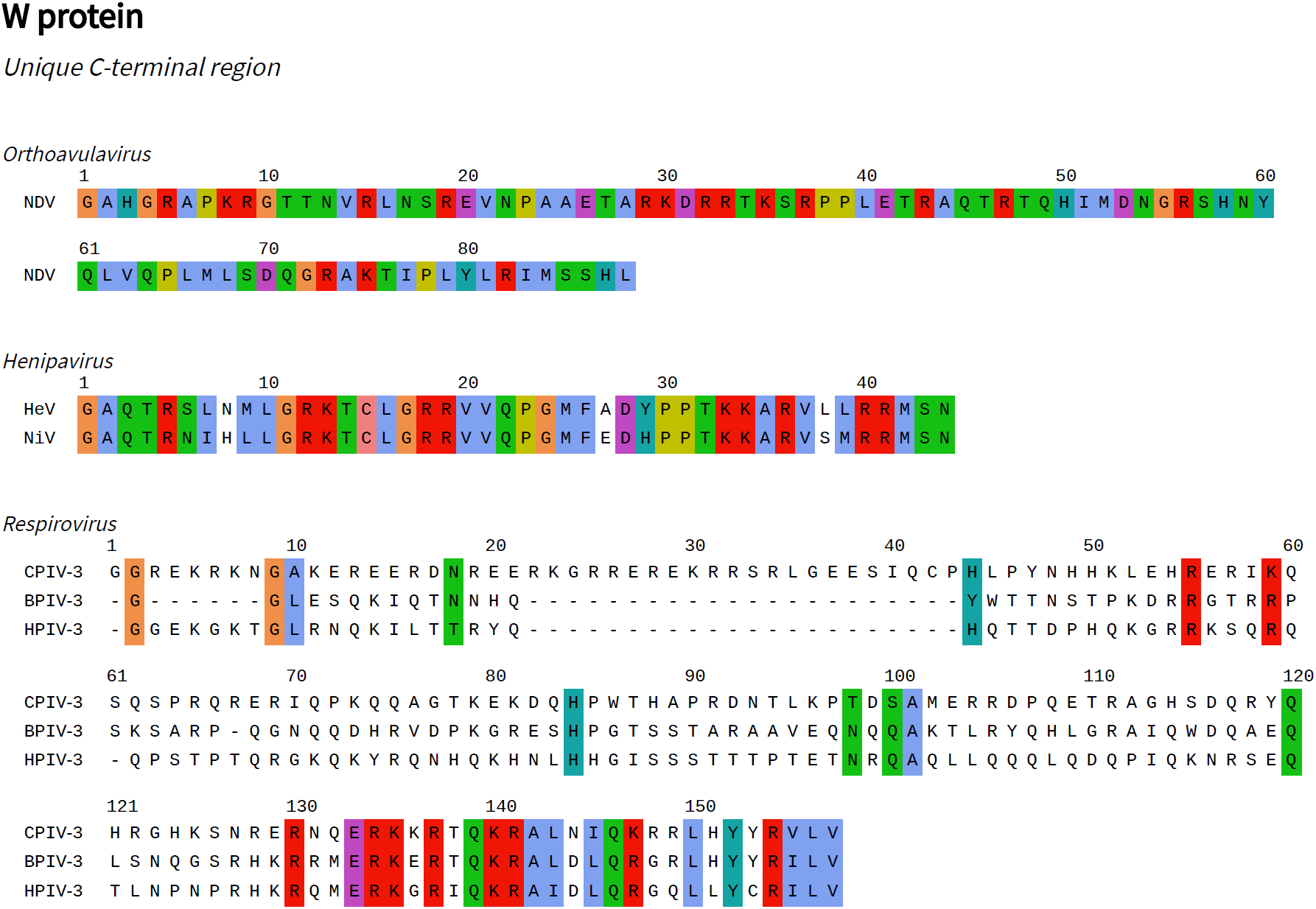
W protein C-terminal regions. For the displayed sequences there is experimental data regarding the cellular localisation or function of the W protein, or a W protein homolog in another virus. All numbering is relative to the start of the unique C-terminal region. Sites are coloured by amino acid characteristic if the characteristic is 100% conserved at the alignment position. Under the ClustalX colouring scheme hydrophobic residues are blue, positively charged residues – red, negatively charged residues – magenta, polar residues – green, cysteine – pink, glycine – orange, proline – yellow, and aromatic residues – cyan (Larkin et al., 2007).

For HPIV-3, in an early study, joint interruption of the V and W open reading frames attenuated viral replication (although individual interruptions had no effect; Durbin et al. (1999)). In interpreting this result, it should be noted that the V protein of HPIV-3 is abnormal, and likely to be expressed in truncated form (Section 3.2). A more recent study also suggests that the C-terminal region of the W protein promotes viral transcription and replication, and is potentially also involved in the downregulation of beta interferon expression (Roth et al., 2013). The C-terminal regions of HPIV-3, bovine parainfluenza virus 3 (BPIV-3; genus: *Respirovirus*), and caprine parainfluenza virus 3 (CPIV-3; genus: *Respirovirus*) W proteins have strong sequence similarity which is itself suggestive of shared function (**Figure 2**).

For remaining paramyxoviruses, the unique region of W may not necessarily possess any biological function at all, and is often very short (2 aa in SeV, 6 aa in MeV, 11 aa in MuV; Chinnakannan et al. (2014); Horikami et al. (1996); Curran et al. (1991); Paterson and Lamb (1990)). However the W protein could still potentially exert biological effects through its shared N-terminal region, with synthesis of W potentially being more rapid than the synthesis of either P or V.

## 3 Phylogeny of cotranscriptional editing

Let *p*(*G*_*m*_) be the empirical probability of the viral transcription machinery inserting *m* guanosines at the mRNA edit site. The most direct source of information on the nature of this probability distribution (i.e. the relative abundances of the transcripts) comes from sequencing the mRNA produced in virally infected cells. However, as Wignall-Fleming et al. (2019) have highlighted, if mRNA preparations are contaminated with anti-genomic RNA, the results may not faithfully reflect the actual abundance of mRNA, and the occurrence of the unedited transcript could be overestimated. Furthermore, several studies have noted that transcript abundance varies with time post-infection (Kulkarni et al., 2009; Qiu et al., 2016). In both cases the proportion of V and W transcripts increased as the infection progressed, though neither the mechanism nor functional implications are understood. Finally, it is noted that while mRNA abundances are often assumed to be related to encoded protein abundances, this may not hold in practice (Liu et al., 2016).

With these caveats noted, the experimentally derived probability distributions (or *edit patterns*) for 26 paramyxoviruses are displayed in **Figure 3**. The maximum observed insert size was *G*_14_ in NiV (Lo et al., 2009). Additional data on mRNA abundance, not displayed in the figure, can be found in the following publications – SeV: Pelet et al. (1991); Kato et al. (1997); NiV: Kulkarni et al. (2009); MeV: Liston and Briedis (1994); Millar et al. (2016); Donohue et al. (2019); NDV: Mebatsion et al. (2001); Yang et al. (2019); BeiV: Audsley et al. (2016b), TevPV: Johnson et al. (2019); Burroughs et al. (2015); HPIV-2: Ohgimoto et al. (1990); MuV: Takeuchi et al. (1990); CeMV: Bolt et al. (1995); PPRV: Mahapatra et al. (2003); PDV: Blixenkrone-Möller et al. (1992); PorPV: Berg et al. (1992).

**Figure 3:**
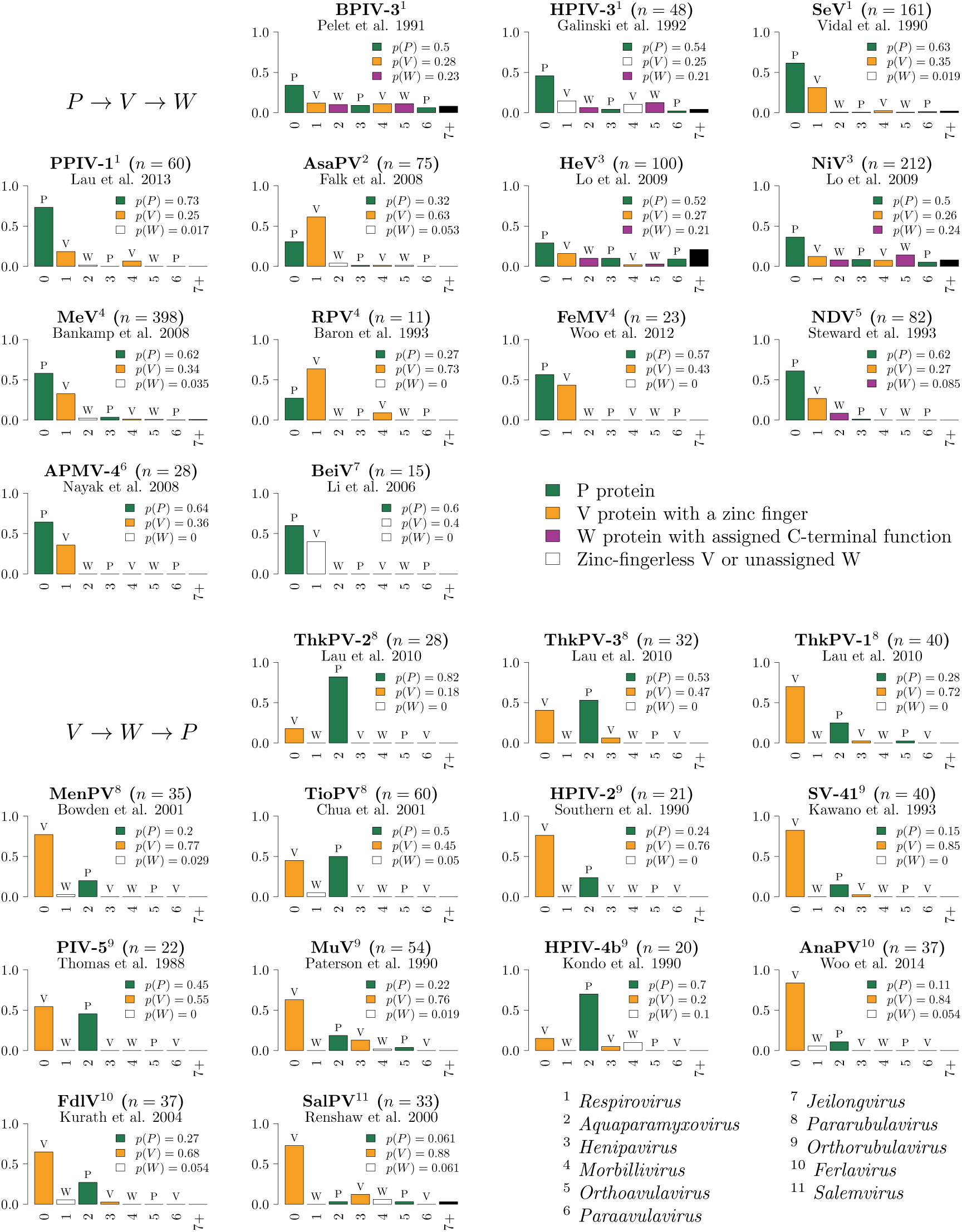
Empirical probability distributions (edit patterns) describing guanosine nucleotide insertion at the P gene edit site. The total proportion of transcripts which encode the three functionally distinct mRNA species are indicated for each experiment. The bulk of the experimental data was obtained by cDNA sequencing, for which the number of sequenced transcripts *n* is specified. Experimental data for BPIV-3 was obtained by a primer extension method acting directly on the mRNA population. Viral genera indicated in bottom right, see Section 6 for virus names.

Fundamental differences that exist between these systems reflect evolutionary events which have occurred throughout the history of the family. The following events are minimally required to explain the data: i) gain of the editing system, ii) loss of the editing system, iii) evolution of the V protein zinc finger motif and gain of biological function, iv) loss of the V protein zinc finger and associated function, v) switching of the edit mode and adaption of the edit pattern, and vi) acquisition of unique function by the W protein. We estimated the evolutionary history of the *Paramyxoviridae* and inferred the ancestral lineages where these events occurred as follows: for each event we imputed the occurrence of the event onto branches such that the number of events required to explain the states observed at the leaves in the tree is minimised (**Figure 4**). This is the maximum parsimony model. A limitation of this model is that it does not account for the full functional diversity of the V protein, which has multiple biological activities (Section 2.2).

**Figure 4:**
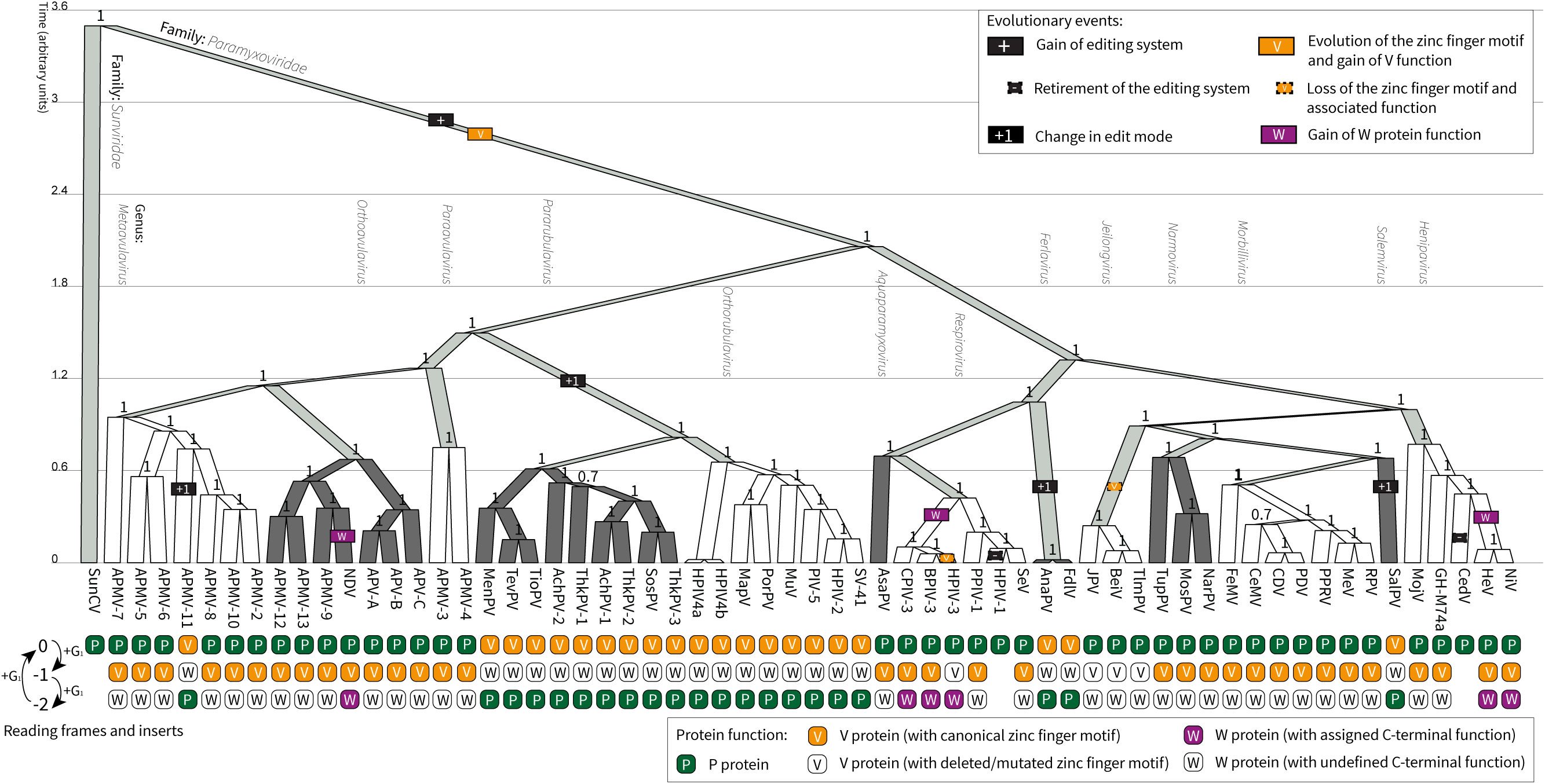
Phylogeny of the *Paramyxoviridae*. Tree created from an alignment of the viral L protein, with Sunshine coast virus (SunCV; Hyndman et al. (2012)) as an outgroup. Coloured rectangles on branches indicate a hypothesised evolutionary event occurring some time in that lineage. Clade posterior supports are shown on the internal nodes. Branches lengths are proportional to time such that there is an average of 1 amino acid substitution per unit of time. Clades are alternatingly coloured by viral genera. See Section 6 for virus names. Tree visualised using UglyTrees (Douglas, 2020).

### 3.1 The editing system and evolution of the V protein

This editing system has not been detected beyond the *Paramyxoviridae* (Hyndman et al., 2012). Therefore, the P gene editing system likely came into existence only once – in the lineage which led to the *Paramyxoviridae*. This event was coupled with the origin of the V protein; the evolution of its unique zinc binding motif; and the gain of many of its conserved functions (**Figure 4**). However the timing of these events cannot be resolved.

### 3.2 Partial or complete loss of the V protein

Under a maximum parsimony model, the V protein has been lost entirely on two independent occasions, both associated with the loss of the editing system (**Figure 4**). The C-terminal zinc binding domain has also been deleted, or significantly mutated, on two further occasions.

Loss of the V protein is associated with retirement of the cotranscriptional editing system. This occurred in both the lineage which lead to Human parainfluenza virus 1 (HPIV-1; genus: *Respirovirus*) and in the lineage which lead to Cedar virus (CedV; genus: *Henipavirus*). As these viruses once employed the 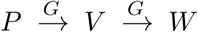 mode, loss of the editing system was axiomatically coupled with loss of both V and W protein expression. It is possible that loss of V protein activity preceded loss of the edit system, but this cannot be determined. Retirement of the editing system appears impossible for viruses employing the 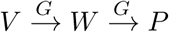 edit mode because the P protein is essential for polymerase function.

In both HPIV-1 and CedV, the edit site is not identifiable using sequence analysis and edited transcripts could not be detected experimentally (Matsuoka et al., 1991; Marsh et al., 2012). In HPIV-1, the conserved V protein sequence is apparent in the genome however there is no clear mechanism for protein production due to the presence of multiple stop codons in the reading frame (Matsuoka et al. (1991); **Figure 5**). This suggests that loss of V occurred quite recently in evolutionary history and there has been insufficient time for the sequences to diverge, creating a pseudogene. For CedV, the V protein sequence is undetectable (Marsh et al., 2012).

**Figure 5:**
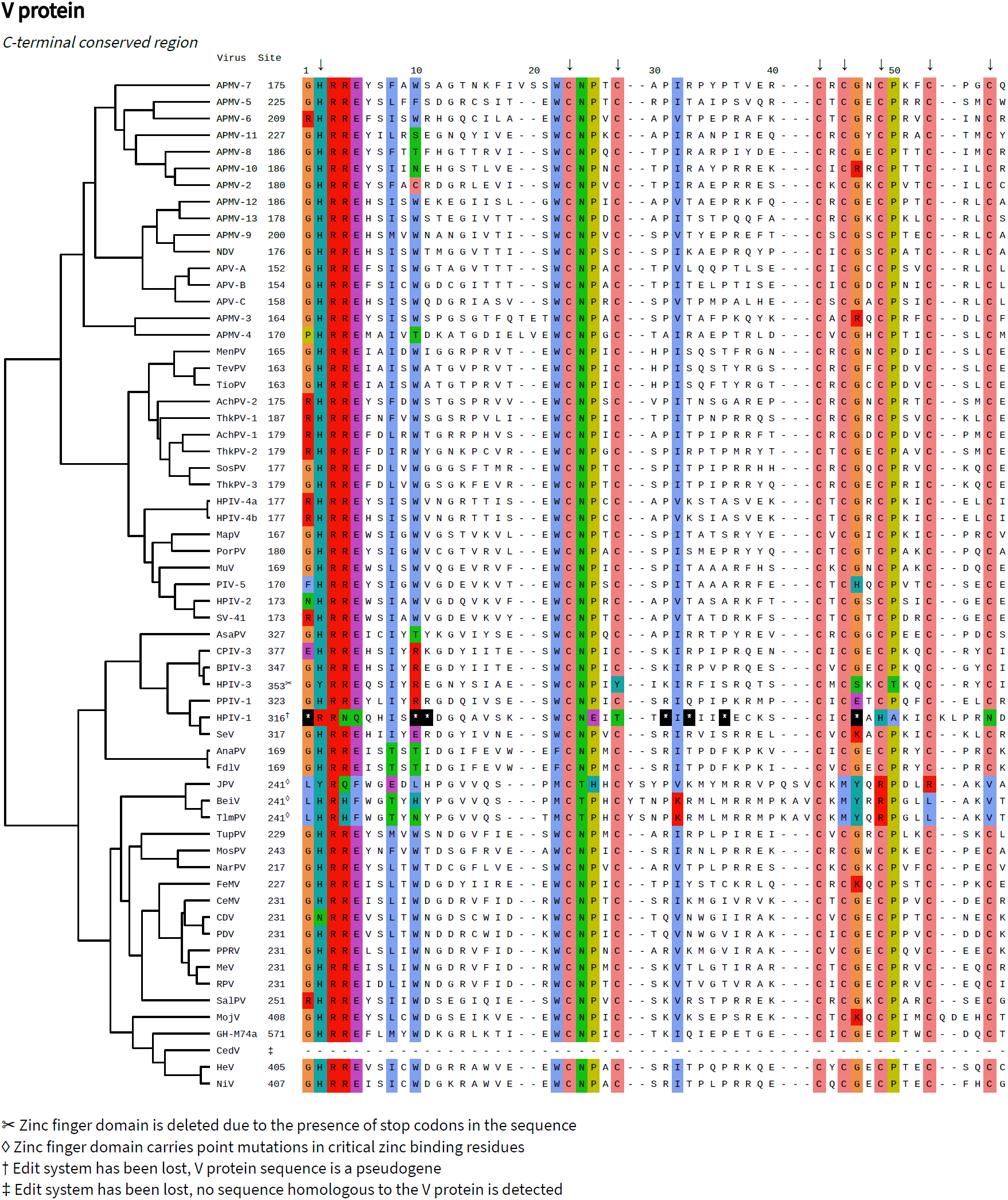
Cysteine-rich C-terminal regions of the V protein. The first amino acid in each aligned sequence is numbered, relative to the start of the V protein. The arrows at the top of the alignment indicate residues whose side chains directly coordinate bound zinc ions, according to a structural study on the PIV-5 V protein (Li et al., 2006a). Asterisks denote stop codons. Sites are coloured by amino acid group if a group is at least 70% conserved at the alignment position (colour scheme indicated in **Figure 2**). The tree is the same as that in **Figure 4**.

In the case of HPIV-3, the edit site is operational (Galinski et al., 1992) and the zinc finger motif is detectable in the genome by sequence analysis (**Figure 5**). However, several stop codons between the edit site and the zinc finger prohibit production of the full-length V protein (unless further non-canonical transcriptional or translational mechanisms are invoked (Galinski et al., 1992)). There are also two mutations in positions which are directly involved in zinc coordination (**Figure 5**). This suggests the V protein zinc finger is a pseudogene, similar to the situation in HPIV-1. In protein-based analysis of infected cells, the full V protein was not detected but a truncated variant which lacks the conserved C-terminal region was (Roth et al., 2013). Overall, current evidence suggests that the V protein of HPIV-3 is expressed in a truncated form lacking the canonical zinc binding motif. Its functional status is unclear.

Finally, in the case of the *Jeilongviruses*, the V protein C-terminal domain has been retained, but with mutation of several critical residues involved in zinc coordination (**Figure 5**). The C-terminal region does not interact with STAT1 or STAT2 (Audsley et al., 2016b), which is a conserved function of other paramyxoviral V proteins (Puri et al., 2009; Röthlisberger et al., 2010). Nonetheless, the Jeilongviral V protein has retained other functions, such as the ability to interact with MDA5 (Audsley et al., 2016b). This finding in particular highlights the multi-functional nature of the V protein, and the limitations of a nomenclature in which its multiple functionalities are not fully explicated.

### 3.3 Switching of edit modes and adaption of edit patterns

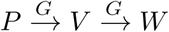 was likely the edit mode of the last common ancestor of the *Paramyxoviridae*.

Under a maximum parsimony model, the editing system has switched to the 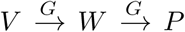 mode four times during evolutionary history (**Figure 4**). These events occurred in the lineages that lead to: 1) Avian paramyxovirus 11 (APMV-11; genus: *Metaavulavirus*), 2) the *Rubulavirinae* subfamily, 3) the *Ferlaviruses*, and 4) Salem virus (SalPV; genus: *Salemvirus*). Edit patterns have been experimentally investigated for three of these four clades: 10 rubulaviruses (Lau et al., 2010; Bowden et al., 2001; Chua et al., 2001; Southern et al., 1990; Ohgimoto et al., 1990; Kawano et al., 1993; Thomas et al., 1988; Paterson and Lamb, 1990; Takeuchi et al., 1990; Kondo et al., 1990), 2 *Ferlaviruses* (Woo et al., 2014; Kurath et al., 2004), and SalPV (Renshaw et al., 2000).

In general the edit patterns of viruses that retain the ancestral 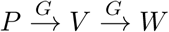 edit mode (**Figure 3**, top panel) are quite different to those of viruses that have subsequently adopted the 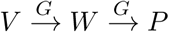 edit mode (**Figure 3**, bottom panel). In the former, *G*_0_ and *G*_1_ insertions are most frequently observed, while in the latter, *G*_0_ and *G*_2_ insertions predominate. It seems clear that edit patterns have co-evolved with edit modes to maintain adequate production of P and V transcripts. In two clades (within the *Respirovirus* and *Henipavirus* genera), the edit patterns are long-tailed, and a significant fraction of the transcripts have more than 2 guanosine nucleotides inserted.

The edit pattern of SalPV (**Figure 3**, bottom panel) appears to be an outlier (Renshaw et al., 2000). *The G*_0_-centric distribution resembles those of viruses using the 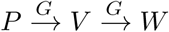 mode, and the relative abundance of P transcripts is very low. Given the taxonomic position of SalPV, as the most immediate outgroup of the *Morbilliviruses* (**Figure 4**), it could be that this is a virus that has switched edit mode but not yet adaptively evolved the edit pattern.

### 3.4 Acquisition of unique function by the W protein

Under our model, the W protein has evolved a novel function associated with its unique C-terminal region on 3 independent occasions (**Figures 2 and 4**): once for NDV (Yang et al., 2019; Karsunke et al., 2019), once for the henipaviral clade comprised of HeV and NiV (Shaw et al., 2005; Lo et al., 2009; Edwards et al., 2020), and once for the respiroviral clade composed of BPIV-3, HPIV-3, and CPIV-3 (Durbin et al., 1999; Pelet et al., 1991). There are varying levels of experimental evidence supporting the existence of a W protein function in these three clades (see Section 2.3). For the remaining paramyxoviruses, W has no known function. Rather, it is more likely that the expression of W is an inevitable byproduct of the editing system; an evolutionary spandrel (Gould and Lewontin, 1979).

For the most part, W transcripts are produced quite rarely (**Figure 3**). However, this does not appear to be the case for two clades where W has acquired function. Instead, the edit pattern is long-tailed, and the probability of producing a W transcript 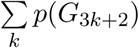 ranges from 21 to 24% in HeV, NiV, BPIV-3, and HPIV-3 (Lo et al., 2009; Pelet et al., 1991; Galinski et al., 1992), and sometimes even higher in temporal analyses (Kulkarni et al., 2009).

In contrast, production of W is not significantly elevated for NDV (Steward et al., 1993; Mebatsion et al., 2001). The overall proportion of W transcript in NDV is estimated at around 8-9% (Steward et al., 1993; Qiu et al., 2016; Yang et al., 2019) or as low as 2.4% (Mebatsion et al., 2001). However, experiments studying the effects of W protein knockout on viral replication (Yang et al., 2019), suggest that these low transcript abundances are optimal for fulfilling the unknown biological function of the NDV W protein (Section 2.3).

## 4 Molecular mechanism of cotranscriptional editing

In the *Paramyxoviridae*, cotranscriptional editing results from transcriptional slippage. This same process facilitates overprinting in other viral families including the *Filoviridae* (Sanchez et al., 1996; Shabman et al., 2014) and *Potyviridae* (Olspert et al., 2015), as well as various prokaryotes (Larsen *et al*., 2000; Penno *et al.*, 2015). Slippage sites can also rescue an organism from deleterious frameshift mutations (Tamas et al., 2008).

Transcription has been extensively studied, recently at the single-molecule level for the RdRP of bacteriophage *ϕ*6 (Dulin et al., 2015a,b) and DNA-dependent RNA polymerases of prokaryotes, eukaryotes, and DNA viruses (Abbondanzieri et al., 2005; Shaevitz et al., 2003; Dangkulwanich et al., 2013; Larson et al., 2012; Skinner et al., 2004; Douglas et al., 2020). These studies have provided significant insights into the mechanisms underlying transcription elongation.

In this final section, we discuss cotranscriptional editing in the *Paramyxoviridae* under the framework presented in the single-molecule literature, noting some additional complexities which arise from the viral genome being packaged within a nucleocapsid.

### 4.1 Transcription elongation by RNA polymerases and slippage

Under a simple Brownian ratchet model, transcription elongation can be modelled as a cycle involving three canonical steps (Bar-Nahum et al. (2005); Abbondanzieri et al. (2005); **Figure 6**, large arrows). First, RNA polymerase steps forward along the template from the pretranslocated to the posttranslocated state, which frees the enzyme’s active site. Second, a complementary nucleoside triphosphate (NTP) binds to the active site. Third, the bound NTP is incorporated onto the 3*1* end of the mRNA and pyrophosphate is released, thus restoring the system to the pretranslocated state.

**Figure 6:**
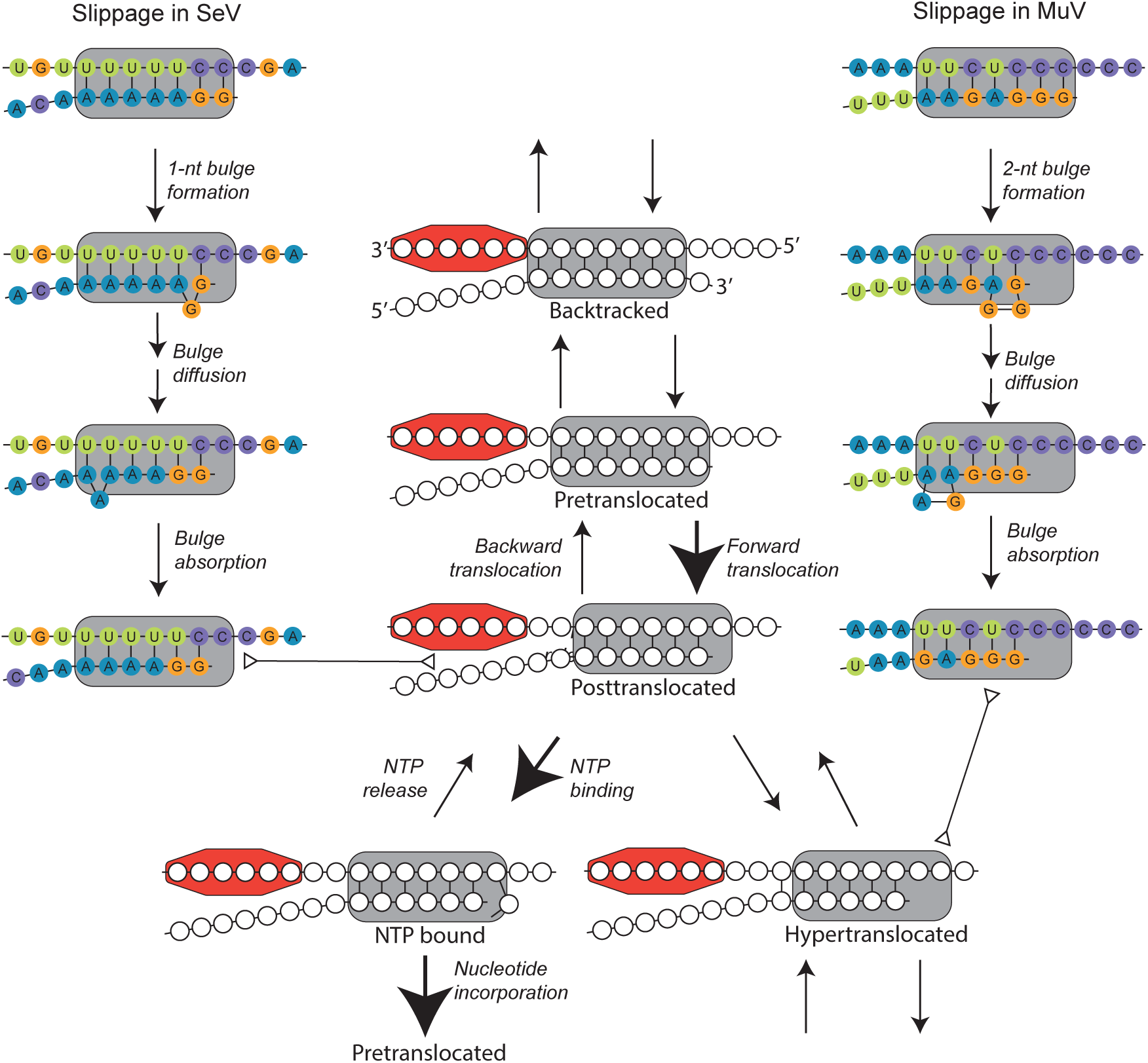
State diagrams of Brownian ratchet and slippage models. Plausible stuttering pathways for SeV (accession: AB039658; genomic position: 2783) and MuV (accession: EU884413; genomic position: 2432) are shown, with a RNA/mRNA hybrid of 7 bp in length. A nucleoprotein protomer bound to the viral genome (top strand) is depicted by the coloured octagon. Large arrows indicate the canonical transcription elongation pathway, double-ended triangular arrows denote equivalency between two connecting states, and unlabelled arrows describe translocation reactions. While slippage initialises in the pretranslocated state in this diagram, the actual state where this process initialises is unknown.

Through backtracking, where the polymerase translocates upstream along the template (Komissarova and Kashlev, 1997; Abbondanzieri et al., 2005), and hypertranslocation, where it translocates downstream (Yarnell and Roberts, 1999), the polymerase can arrive at a catalytically inactive state (**Figure 6**). These processes can lead to transcriptional pausing (Saba et al., 2018; Artsimovitch and Landick, 2000). In the case of paramyxoviruses, back-tracking and hypertranslocation may be inhibited by the presence of genome-wrapped nucleoproteins acting as “roadblocks”, analogous to the role played by nucleosomes in eukaryotes (Nudler, 2012).

Slippage involves the movement of one sequence in the product/template hybrid relative to the other, which can lead to imperfect basepairing. Slippage was hypothesised by Streisinger et al. (1966) as one of the primary mechanisms of indel events. The mechanism is thought to involve formation of a nucleotide bulge near the 3*1* end of the mRNA (Garcia-Diaz and Kunkel, 2006). If the bulge forms in the nascent strand, an insertion can result, whereas a bulge in the template strand can lead to a deletion.

By applying varying forces to individual dsDNA molecules, Kühner et al. (2007) and Neher and Gerland (2004) hypothesise that slippage occurs in three steps (**Figure 6**). First, a bulge forms on one side of the hybrid. This initial reaction must overcome a large Gibbs energy barrier. Second, the bulge diffuses along the hybrid. Diffusion is likely to be quite rapid (Woodson and Crothers, 1987), and favoured if Watson-Crick basepairing is maintained in the bulged hybrid. Third, the bulge is absorbed at the other end of the hybrid. During transcription, the next nucleotide could be incorporated before or after absorption, if absorption even occurs (Tippin et al., 2004).

In principle, transcriptional slippage could be initialised from any one of the states available to the polymerase (backtracked, pretranslocated, posttranslocated, or hypertranslocated), and it is not known if the the process of bulge formation is linked to the translocation state.

### 4.2 Transcriptional slippage in paramyxoviruses

Through transcriptional slippage, a single templated nucleotide can be copied multiple times (stuttering). Stuttering is the apparent mechanism of cotranscriptional editing in paramyxoviruses and can explain many of the edit patterns presented in **Figure 3**.

*-*The two distinct modes of editing (i.e. 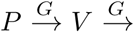 and 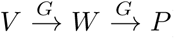)are encoded by quite different sequences (**Figure 7**).

**Figure 7:**
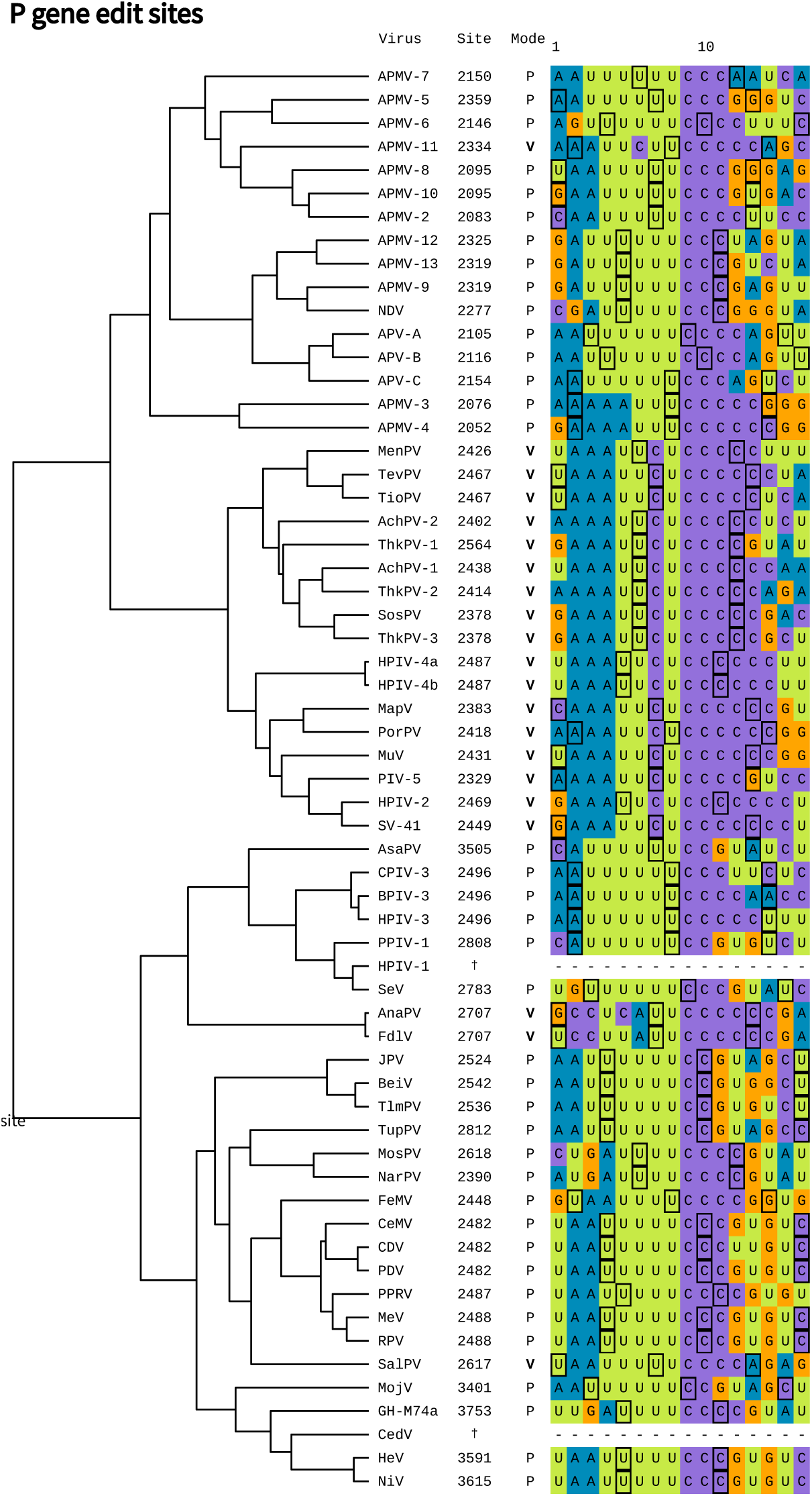
Edit site sequences in the paramyxoviruses. The sequences of the negativesense (genomic) RNA are displayed. The numbers indicate the genomic position of the first displayed nucleotide. 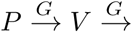 and 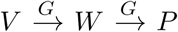 edit modes are abbreviated to P and **V** respectively. Nucleoprotein phases are displayed; the first nucleotide within each nucleoprotein protomer is highlighted in black. This tree is the same as that in **Figure 4**.

The edit sites among viruses which employ the 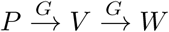 edit mode are conserved. Using the PROSITE notation (Sigrist et al., 2002), this (genomic-sense) edit site motif can be described by U(3,6)–C(2,6). In SeV, the edit site sequence is transcribed into AAAAAGgG, where the lower-case g is the stutter site i.e. the site reiteratively transcribed from the template (Hausmann et al., 1999b,a; Vidal et al., 1990b). Under the stuttering model, inserts are added as follows (**Figure 6**, left hand side): 1) a bulge forms in the 3*1* mRNA of the RNA/mRNA hybrid. 2) The bulge is free to diffuse along the hybrid. Although the bulge is thermodynamically disfavoured, it can occur because of U/A and non-canonical U/G basepairing which are maintained throughout diffusion. 3) In no particular order, the bulge is absorbed at the 5*1* end and the lower-case g can be transcribed again. Each iteration of these three steps is associated with a *G*_1_ insertion.

In contrast, the edit sites across the four clades of the 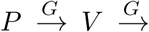 group are quite distinct from one another. SalPV is anomalous, and its edit site sequence resembles the 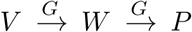 group (Renshaw et al., 2000). This could explain the relatively low amounts of P transcript produced (**Figure 3**). The *Ferlavirus* edit site is distinct from all other known edit sites (Woo et al., 2014; Kurath et al., 2004) and the mechanism of guanosine insertion is not clear. Through convergent evolution, APMV-11 and the *Rubulavirinae* subfamily have similar edit sites (PROSITE: A(3,4)–U(2)–C–U(1,2)–C(4,7); genomic-sense). In the case of MuV, the edit site (transcribed into UUUAAGAGGG) has been well characterised (Vidal et al., 1990b). Stuttering occurs in a similar fashion to SeV, however the edit sequence allows *G*_2_ inserts (encoding the P protein) to occur at a greater frequency than *G*_1_ inserts (encoding the W protein) due to the formation of a 2 nucleotide bulge (**Figure 6**, right hand side).

Slight variation in the edit site sequence perturbs stuttering of the viral RdRP. For instance, when the length of the poly(A) sequence at the SeV edit site was increased, from A(3)–G(6) to A(8)–G(1), the average number of inserts increased dramatically (Hausmann et al., 1999a). Similarly, when the SeV edit site sequence was mutated to resemble that of BPIV-3, its edit pattern changed correspondingly (Hausmann et al., 1999b). While slippage patterns can also be dependent on cellular environment (Donohue et al., 2019), these results speak to the primary importance of the genome sequence in governing polymerase stuttering.

The roles that nucleoprotein displacement and the rule of six play during cotranscriptional editing have been investigated (Iseni et al., 2002; Hausmann et al., 1996; Kolakofsky, 2016). Changing the nucleoprotein phase around the edit site sequence (of SeV) resulted in an apparent change in edit pattern (Iseni et al., 2002). We computed the expected nucleoprotein phase at the edit site of each virus under the rule of six model. Although nucleoprotein displacement may play a role in editing, the nucleoprotein phase at the edit site does not appear to be well conserved (**Figure 7**).

## 5 Conclusion

In this review, we compiled the genomic sequences of paramyxovirus edit sites (**Figure 7**) and, where available, their experimentally determined edit patterns (histograms of insert sizes; **Figure 3**). We estimated the evolutionary history of editing in the family (**Figure 4**). This analysis suggests that, among the characterised paramyxoviruses, the editing system has independently switched from the 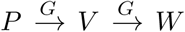 to the 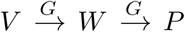 edit mode four times; the V protein zinc finger domain was deleted or significantly mutated twice, the W protein has evolved a known function in its unique region three times; and the cotranscriptional editing system has been lost entirely twice, leading to complete loss of both V and W expression. Although transcriptional slippage provides the mechanism for non-templated base insertion, it is currently unclear how this process is coordinated with either canonical or non-canonical steps of the elongation pathway.

## 6 Virus abbreviations

AchPV 1-2: Achimota viruses 1-2
APMV 2-13: Avian paramyxoviruses 2-13
AsaPV: Atlantic salmon paramyxovirus
BeiV: Beilong virus
CDV: Canine distemper virus
CeMV: Cetacean morbillivirus
FdlV: Fer de Lance virus
HeV: Hendra virus
JPV: J-virus
MeV: Measles virus
MosPV: Mossman virus
MuV: Mumps virus
NDV: Newcastle disease virus
PDV: Phocine distemper virus
PorPV: Porcine rubulavirus
PPRV: Peste-des-petits-ruminants virus
SalPV: Salem virus
SosPV: Sosuga virus
SV-41: Simian virus 41
ThkPV 1-3: Tuhoko viruses 1-3
TlmPV: Tailam virus
AnaPV: Anaconda paramyxovirus
APV A-C: Antarctic penguin viruses A-C
GH-M74a: Ghanaian bat henipavirus
BPIV-3: Bovine parainfluenza virus 3
CedV: Cedar virus
CPIV-3: Caprine parainfluenza virus 3
FeMV: Feline morbillivirus
HPIV 1-4: Human parainfluenza viruses 1-4
MenPV: Menangle virus
MojV: Mojiang virus
MapV: Mapuera virus
NarPV: Nariva virus
NiV: Nipah virus
PIV-5: Parainfluenza virus 5
PPIV-1: Porcine parainfluenza virus 1
RPV: Rinderpest virus
SeV: Sendai virus
SunCV: Sunshine coast virus
TevPV: Teviot virus
TioPV: Tioman virus
TupPV: Tupaia virus

## 7 Algorithms and data availability

Sequences were aligned by M-Coffee (Wallace, 2006) and treated with subsequent manual adjustment using AliView (Larsson, 2014). Phylogenetic tree built with BEAST 2 (Bouckaert et al., 2019) from an alignment of the L protein, and a relaxed clock model Drummond et al. (2006). P/V/W sequences, L alignment, and BEAST 2 input/output files are available at https://github.com/jordandouglas/ParamyxovirusSlippageEvolution.

